# Cell wall-mediated maternal control of apical-basal patterning of the kelp *Undaria pinnatifida*

**DOI:** 10.1101/2024.04.26.591308

**Authors:** Eloise Dries, Yannick Meyers, Daniel Liesner, Floriele Gonzaga, Jakob Becker, Eliane E. Zakka, Tom Beeckman, Susana M Coelho, Olivier De Clerck, Kenny A Bogaert

## Abstract

The role of maternal tissue in the control of embryogenesis remains enigmatic in many complex organisms. Here, we investigate the contribution of maternal tissue to apical-basal patterning in the kelp embryo. Using a modified kelp fertilisation protocol which yields synchronously developing kelp embryos, we show that detachment from maternal tissue leads to compromised robustness of apical-basal patterning. Detached embryos are rounder and often show aberrant morphologies. Furthermore, absence of contact with maternal tissue increases parthenogenesis, highlighting the critical role of maternal signals in the initial stages of kelp development. When zygotes are detached from the female gametophyte while part of the oogonial cell wall still remains attached to the egg, the proper apical-basal patterning is rescued showing a key role for the connection to the maternal cell wall in developmental patterning in kelps. This observation is reminiscent of another brown alga, *Fucus*, where the cell wall has been shown to play a key role in cell fate determination. In the case of kelps, the maternal oogonium mediates basal cell fate determination by providing an extrinsic patterning cue in its extracellular matrix to the future embryo. Our findings suggest a conserved mechanism across phylogenetically distant oogamous brown algal lineages, where localised secretion of sulphated F2 fucans mediate establishment of the apical-basal polarity.

## Introduction

In many complex multicellular organisms such as animals, red algae and land plants, zygotes develop in close association with the maternal tissue (Borg et al., 2023; Kölle et al., 2020; Woudenberg et al., 2024). In plants, embryo development typically begins by establishing apical-basal polarity and an initial asymmetric division of the zygote which is embedded within maternal tissue. The extent of guidance, protection and developmental robustness that the maternal tissue can provide to plant embryogenesis remains an open question (Woudenberg et al., 2024).

Contrary to land plants and red algae, most brown algal species release their gametes in the environment. Zygotes therefore develop free from maternal tissue. Additionally, development of the zygotes may occur naturally synchronised by simultaneous gamete release and fertilisation. Consequently, studies on brown algae have offered important insights in cell polarization and embryonic patterning in a lineage that evolved multicellularity independently from plants (reviewed in: Bogaert et al., 2022; Coelho & Cock, 2020; De Smet & Beeckman, 2011). Specifically, research on the brown alga *Fucus* has highlighted the critical role of cell fate determinants present in the extracellular matrix (ECM) (Berger et al., 1994; Bouget et al., 1998).

The role of maternal tissue in early development of brown seaweeds displays considerable variation. In *Fucus*, eggs are released in the environment and zygotes undergo polarization entirely *de novo*, either in response to light or the entry of paternal sperm (Hable & Kropf, 2000). Conversely, *Dictyota* zygotes retain a maternally determined direction for the polarization vector, while the sense of the vector is determined by the direction of light (Bogaert et al., 2017). In oogamous brown algae of the order Laminariales and family Phyllariaceae (Tilopteridales) (here collectively referred to as ‘kelps’) the release of female gametes is incomplete. Eggs and embryos remain physically connected to the female gametophyte after extrusion from the oogonium. However, relatively little is known on the relevance of this connection to the maternal tissue in apical-basal patterning.

The study of kelp embryogenesis is further conflated by the occurrence of parthenogenetic development (Druehl et al., 2005; tom Dieck 1992), with eggs being able to develop in the absence of fertilisation. Parthenotes typically detach more easily (tom Dieck 1992; Klochkova et al., 2019; Martins et al. 2017) and are rounder (Schreiber 1930; tom Dieck 1992; Le Gall et al. 1996; Fang et al., 1979). This aberrant morphology and tendency to detach are used as indications for parthenogenesis (tom Dieck et al. 1992; Martins et al. 2017; Liboureau et al. 2024).

In kelp, the eggs bear flagella, similar to the male gametes (Motomura, 1990; Motomura & Sakai, 1988). Only recently, it was shown that these female flagella anchor the egg to the oogonium (Klochkova et al., 2019). The occurrence of flagella, basal bodies, flagellar roots and a polarised cytoplasm (Motomura, 1994) imply that the egg is a polarised cell. The anchor flagella are considered essential for the polarity and survival of the sporophyte (Klochkova et al., 2019). Exploring the fate of detached progeny in ecology is crucial for understanding dispersal mechanisms and adaptability of kelp populations to environmental changes.

Here, we used the kelp *Undaria pinnatifida* to test whether apical-basal polarity in kelp embryos is established entirely autonomously or whether signaling from the maternal gametophytic tissue is necessary for correct patterning and early embryonic development. In order to exclude confusion with putative polarity defects of parthenotes, we investigate the events underlying early embryogenesis and contrast these with parthenogenesis.

We show that kelp embryos lacking maternal contact lose their apical-basal polarity, leading to defects in developmental patterning which resemble those observed in partheno-sporophytes. Using time-lapse imaging and micro-dissection, we can distinguish between the impacts of the cell wall and cytoplasmic signals from the maternal tissue. We show that the oogonial cell wall is required for apical-basal patterning of the embryo. Additionally, we show that Golgi-derived cell wall material is deposited in the oogonium, reminiscent of another brown alga, *Fucus*, where the cell wall has been shown to play a key role in cell fate determination. We propose that factor(s) secreted into the oogonial cell wall by the maternal gametophyte determine the fate of the basal cell in the developing embryo, contributing to developmental patterning robustness of the diploid sporophyte.

## Materials and Methods

### Algal strains and growth conditions

Experiments were carried out using unalgal laboratory cultures of respectively *Undaria pinnatifida* (Upin_NL1f (female) and Upin_NL1m (male)), isolated from a single sporophyte sampled in Westerschelde (2020, The Netherlands). Additional observations were made on *Saccorhiza polyschides* (PH-IS_016f (female) and PH-IS_001m (male) (Perharidy, France), *Macrocystis pyrifera* (CVe13f (female) and CVe30m (male) (2006, Curaco de Vélez, Chile) and *Saccharina latissima* (Slat_FR1f (female), Slat_FR1m (male) (2020, Brittany, France) (See supplemental experiment, **Figure S6**). For *Undaria*, fertility was induced by fragmenting male and female gametophytes. *Undaria* cultures were grown separately at 18°C at 80 μmol photons m^−2^·s^−1^ (Lumilux, Cool White, Osram, Germany) and a photoperiod of 12:12 (L:D) in filtered and autoclaved natural seawater enriched with modified Provasoli (mPES) (West and McBride 1999). Fertility of *Macrocystis* and *Saccorhiza* was induced at 12°C under a 16:8 h (L:D) photoperiod with 20-30 µmol photons m^-2^ s^-1^. Fertility for *Saccharina* was induced at 10°C under e 12:12 photoperiod at 10 µmol photons m^-2^ s^-1^.

### Synchronized fertility

To avoid that our control populations contain a mixture of age cohorts released at different days, it is important to control the timing of fertilisation. Therefore we designed a protocol for synchronised kelp fertilisation, making use of two key features of kelp reproduction: in a light-dark cycle, female gametophytes mainly release eggs within 30 minutes of darkness (Lüning, 1981); and a pheromone produced by the released eggs triggers sperm release and attraction (Maier et al., 2001). Filtered medium of female gametophytes containing freshly released eggs collected after 2 hours of darkness was added to fertile male gametophytes to trigger sperm release. After 2 minutes of induction, the male medium containing suspended sperm cells was filtered through a sterile 40 µm nylon mesh to remove the male gametophytes. The sieved suspension with male gametes was combined with the remaining female gametophyte culture to induce fertilisation, followed by incubation in darkness to continue the night cycle. This method ensures the production of a single age cohort of sporophytes by allowing only a short time window for fertilisation.

### Detachment of zygotes

Fertilised suspended female gametophytes were rinsed over a 40 µm nylon mesh to remove loose eggs and parthenotes and zygotes. Fertile gametophytes were pipetted up and down using a Pasteur pipette for 3 minutes to detach zygotes. The culture was then filtered again through a 40 µm nylon mesh and detached zygotes and eggs were isolated in a 24-well plate.

### Parthenogenesis control

For describing and controlling for the effects of parthenogenetic development, we detached unfertilised eggs in female-only cultures. Eggs were detached as described for zygotic development. Parthenogenesis was scored as development beyond a single-cell state at day 4 after detachment.

### Microsurgery in the oogonial cell wall

Using a Sharpoint 4 mm microsurgical knife a precise incision was made between the live gametophyte and the *Undaria* zygote, leaving a piece of the oogonial cell wall attached to the zygote. After four days of cultivation, growth was assessed. In *Saccharina*, detachment of zygotes by pipetting resulted in a mixture of putative zygotes, with some retaining an attached piece of the oogonial cell wall, while others showed no visible remainder of the cell wall. Therefore, microsurgery as described for *Undaria* was not necessary.

### Microscopy

Embryos were photographed at day 1 and 4 with a Zeiss Axioplan 2 light microscope (Carl Zeiss, Jena, Germany) equipped with an AxioCam camera and an Olympus BX-51 microscope (Olympus, Tokyo, Japan) equipped with a digital camera (ToupCamTM, Touptek, China). ImageJ was used to measure several shape characteristics, including area, circularity, roundness, and aspect ratio (AR) (Schneider et al. 2012). Both roundness and circularity values range from 0 to 1, with higher values indicating shapes that closely resemble a perfect circle. Lower circularity values correspond to more irregular shapes, while AR ranges from 1 to positive infinity and describes its elongation or flattening by comparing the major axis to the minor axis.

### Statistics

For comparing two independent groups, when the data were normally distributed and variances were equal as confirmed by Levene’s test, a Student’s t-test was applied. In cases where the data did not meet these assumptions, a non-parametric Wilcoxon rank-sum test was employed. To examine the proportion of embryos to the total count of released eggs in different groups, a generalized linear model (GLM) with a quasi-binomial distribution was utilized to account for the overdispersion in the data. Post hoc comparisons between group levels were conducted using estimated marginal means with Tukey’s adjustment for multiple comparisons to control the family-wise error rate. For comparing the morphometric measurements between groups, controlling for embryo size, a GLM with a Gamma distribution and log link function was used (for AR) or a beta regression model (Cribari-Neto & Zeileis, 2010) with logit link function for circularity and roundness which ranges between 0 and 1 to account for heteroscedasticity. Analyses were conducted in R (version 4.1.2) (R Core Team, 2023).

### Genotyping of embryos and parthenotes

To ascertain that *Undaria* embryos resulted from fertilised eggs, we amplified sex markers M_68_16_2 and M_285_20_2 (Lipinska et al., 2015) using the KAPA3G Plant PCR Kit for direct genotyping. Samples were isolated at 4 days in PCR-grade water, crushed and stored at -20°C. PCRs were conducted with the following PCR conditions (Initial: 95°C for 5min, 40 cycles: 95°C (20sec), 65°C (25sec), 72°C (30 sec), Final: 72°C (7min)).

### Flow cytometry

For the ploidy analysis, small clusters of gametophytes weighing 20-50 mg were dried lightly, finely chopped in 500 μL of nuclei isolation buffer, and treated with 0.5 μg/L Proteinase K, 5‰ SDS, and 0.1 mM Tris HCl at pH 8.0. The nuclei were then filtered through a 10 μm nylon mesh and dyed with 1X SYBR Green I. Following a 10-minute incubation, the DNA content was analyzed using an Agilent NovoCyte Advanteon flow cytometer.

### Toluidine Blue stainings

Cell wall modification in the oogonial cell wall was assayed by Toluidine Blue O staining (TBO), which stains sulfated fucans indicative of polar secretion of Golgi-derived material into the cell wall in *Fucus* (Shaw & Quatrano 1997). Gametophytes were stained for 15 minutes with 0.1 % Toluidine Blue O artificial seawater at pH 1.5. Slides were rinsed three times in 99 % ethanol for approximately 5 minutes and once for 1 hour before being mounted in tap water and photographed.

## Results

### Establishment of apical basal polarity during early development

To describe the timing of the expression of the apical-basal polarization vector, we investigated early embryogenesis in *Undaria.* When female gametophytes reach maturity, eggs are extruded from the oogonium, but remain connected to the oogonial aperture through two flagella (**Figure 1A,B**). Initially spherical, *Undaria* eggs elongate within the first hour after fertilisation (AF) by expansion of the cell volume, and not by mere shape change (**Figure 1A,C, Figure S1, S2**). The basal part of the zygote bulges, and this bulge serves to anchor the embryo to the opening of the oogonial cell wall (**Figure 1, S1**). At approximately 15 hours after fertilisation (AF), a clear hyaline zone at the site of the future cytokinesis plane becomes evident and zygotes undergo a first, asymmetric cell division (**Figure 1D, S1**). The first cell division results in the formation of an apical cell, which will develop into the upper part of the thallus, and a basal cell, which will develop into the lower part of the thallus and secure the embryo to the oogonial cell wall of the gametophyte (**Figure 1A**).

**Figure 1.**
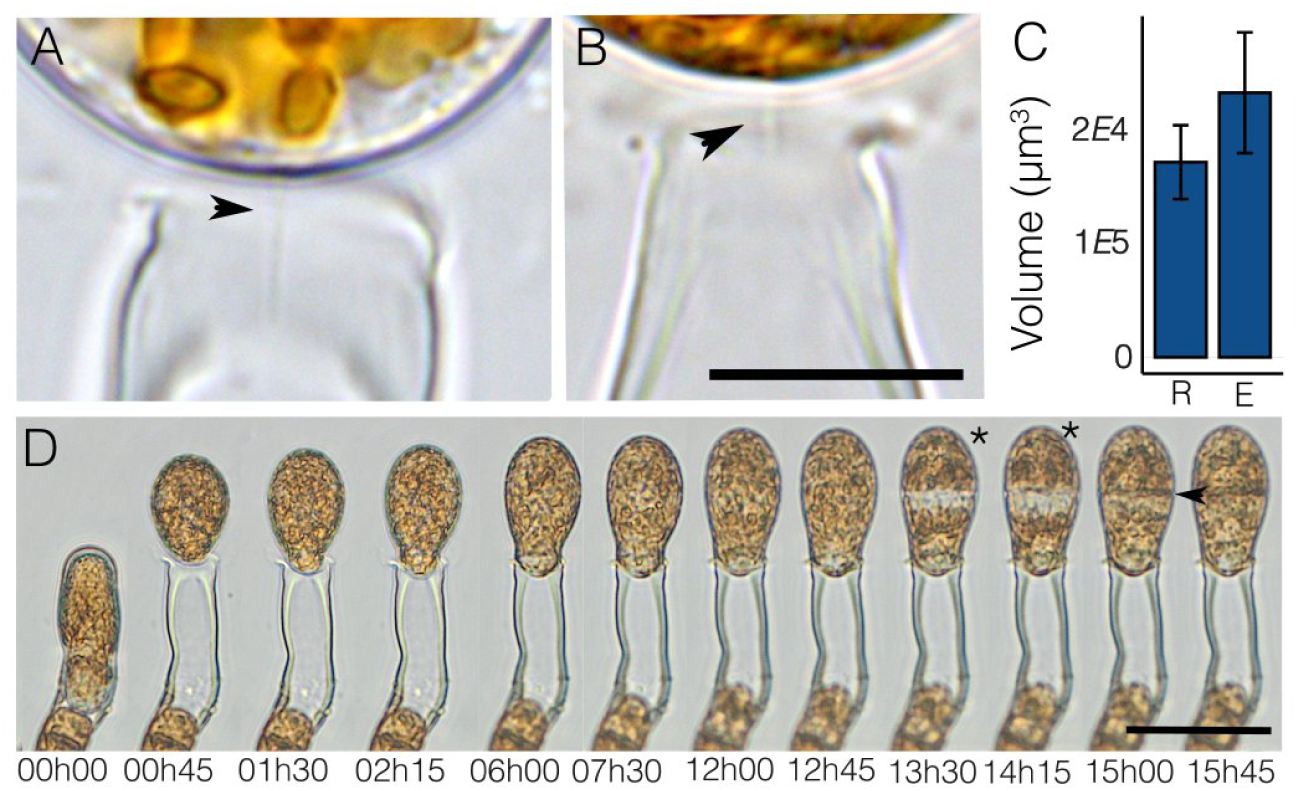
Elongation and first cell division of the *Undaria pinnatifida* zygote. (**A, B**) The embryo is connected to the oogonial cell wall via anchor flagella (arrow head). Scale bar = 5 µm (**C**) Volume increase between round cells (R) and elongated cells (E) (P=0.0006, Wilcoxon rank sum exact test). (**D)** Timelapse of egg release, elongation and first asym metric cell division in the zygote. The first cell division is visible as a hyaline zone (*) before completion of the cell plate. Apical and basal cell have a different fate, the basal cell forms a basal bulge firmly attaching the zygote to the oogonial cell wall. Scale bar = 50 µm.

In order to understand how the apical-basal axis is expressed in normal embryo development, we described development following the first cell division. Zygotes typically divide twice transversely, i.e. perpendicular to the polarization vector, resulting in a 4-celled embryo, (**Figure 2**) resulting in 2 cells derived from the basal cell and 2 cells derived from the apical cell (**Figure 2A,B,C**). The 3 most basal cells divide transversely again, while the most apical cell divides parallel to the apical-basal axis, initiating the process of blade widening (**Figure 2C**). The resulting 2 apical cells divide transversely resulting in a total of 8 segments. These segments further subdivide predominantly parallel to the polarization vector in the apical half. The most basal cells divide substantially less (**Figure 2B,C**). Further growth results in an elongated, one-layered blade with clear apical-basal polarity by day 4 AF (**Figure 2D**). At the same time cells of the female gametophyte gradually fill the empty space created by extrusion of the egg and ultimately make contact with the embryo at approximately day 3 or 4, when the embryo is measuring 40 cells or more (**Figure 2D, S3).** Therefore embryos develop in direct contact only with the oogonial cell wall for the first days.

**Figure 2.**
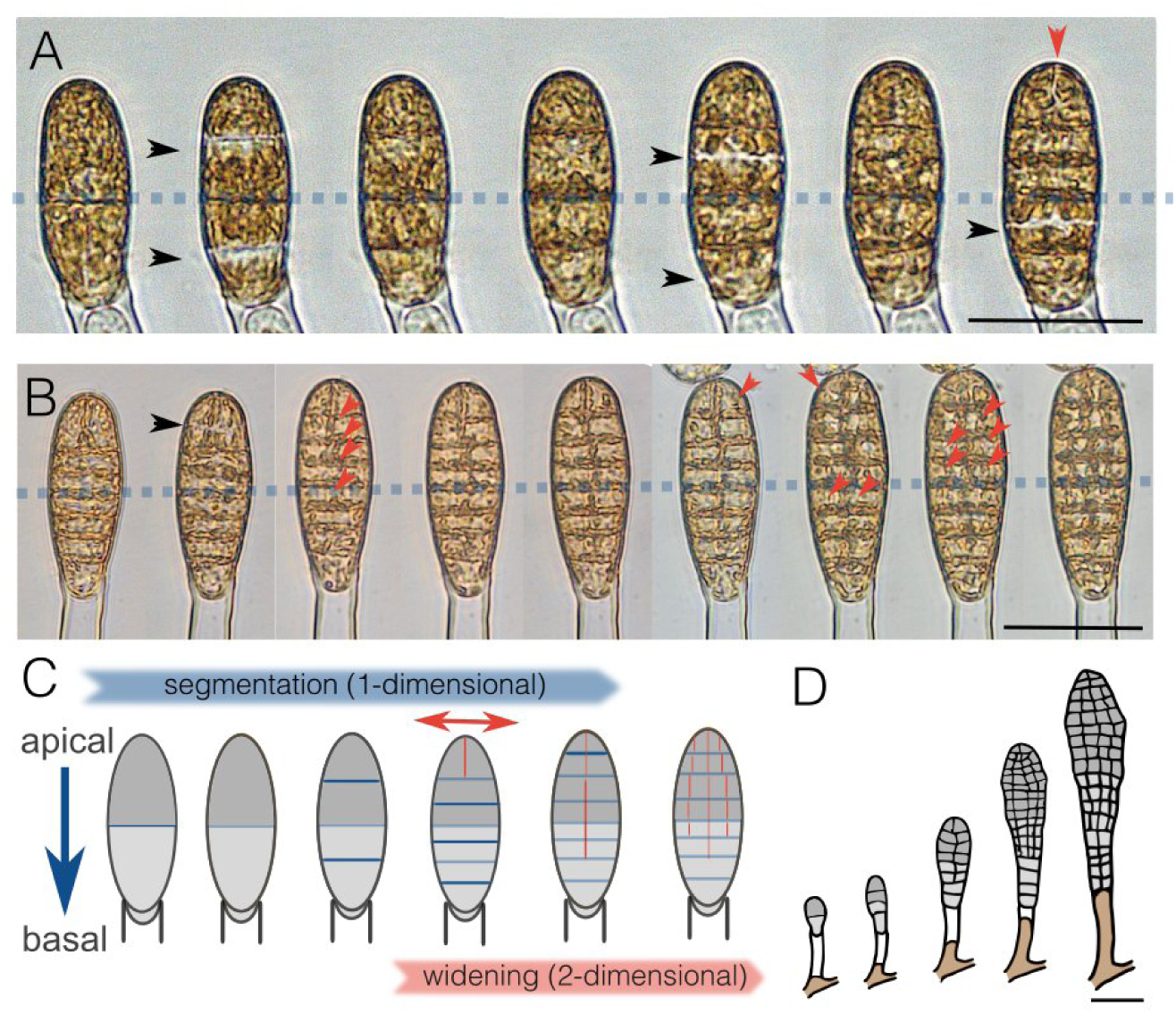
Early development of the *Undaria* embryo. **(A, B**) Both apical and basal cells of the embryo first divide perpendicular to the long axis of the embryo after which cell division changes orientation parallel to the polarization axis (black arrow) in the most apical cell (red arrow). The three most basal cells divide perpendicular again. Blue dashed line separates the apical from the basal cell line. (**C**) Schematic representation of orientation first cell and apical basal gradi ent in cell division. Blue, transversal cell divisions perpendicular on the apical basal vector; red cell divisions, parallel to the apical basal vector. (**D**) Schematic representation of blade expansion at apical end and female gametophyte expand ing into the oogonial tube with new tissue. Drawings are based on time lapse pictures taken at day 0,1,2,3,4. Scale bar = 50 µm.

### Embryos and parthenotes exhibit distinct developmental trajectories

In kelps, eggs may initiate parthenogenetic development in absence of fertilisation (Shan et al. 2013). These parthenotes tend to grow rounder (Ar Gall et al.,1996; Fang et al. 1979), detach more easily (Martins et al., 2017; Klochkova et al., 2019) and consequently an enrichment of parthenogenetic (rounder) individuals in the detached fraction may be expected. Moreover patterning deffects are a sign of parthenogenesis (tom Dieck et al., 1992; Martins et al., 2017; Liboureau et al., 2024). Therefore, in this study, we ensured that any patterning defects are not due to the effects of parthenogenesis. For this reason, we first had to examine the developmental patterns of parthenogenesis to discover how to discern parthenotes from embryos. We examined the developmental patterns of parthenotes in order to distinguish these from proper embryo development and the effect of maternal tissue on apical-basal patterning. To do so, female gametophytes were cultivated in the absence of males for 7 days until they reached fertility and produced eggs. We then scored the presence of parthenotes. We found that parthenogenesis events are rare (1.2 % ± 0.7 %) compared to control populations where eggs remain attached to the maternal gametophyte. We then tested the effect of detaching the eggs and cultivating them in natural SW or in a gametophyte-conditioned medium. The proportion of parthenogenetic embryos dramatically increased when the eggs were artificially detached from the maternal gametophyte (27.6% ± 3%) (**Figure 3C**), regardless of whether gametophyte conditioned medium or fresh medium was applied to the detached eggs. Therefore, it appears that detachment of eggs from the maternal tissue stimulated parthenogenesis, or, in other words, physical contact with the maternal gametophyte prevented initiation of parthenogenesis. Conditioned media does not reverse the effect, suggesting that the physical, continuous presence of the maternal tissue is required to maintain a non-parthenogenetic state.

**Figure 3.**
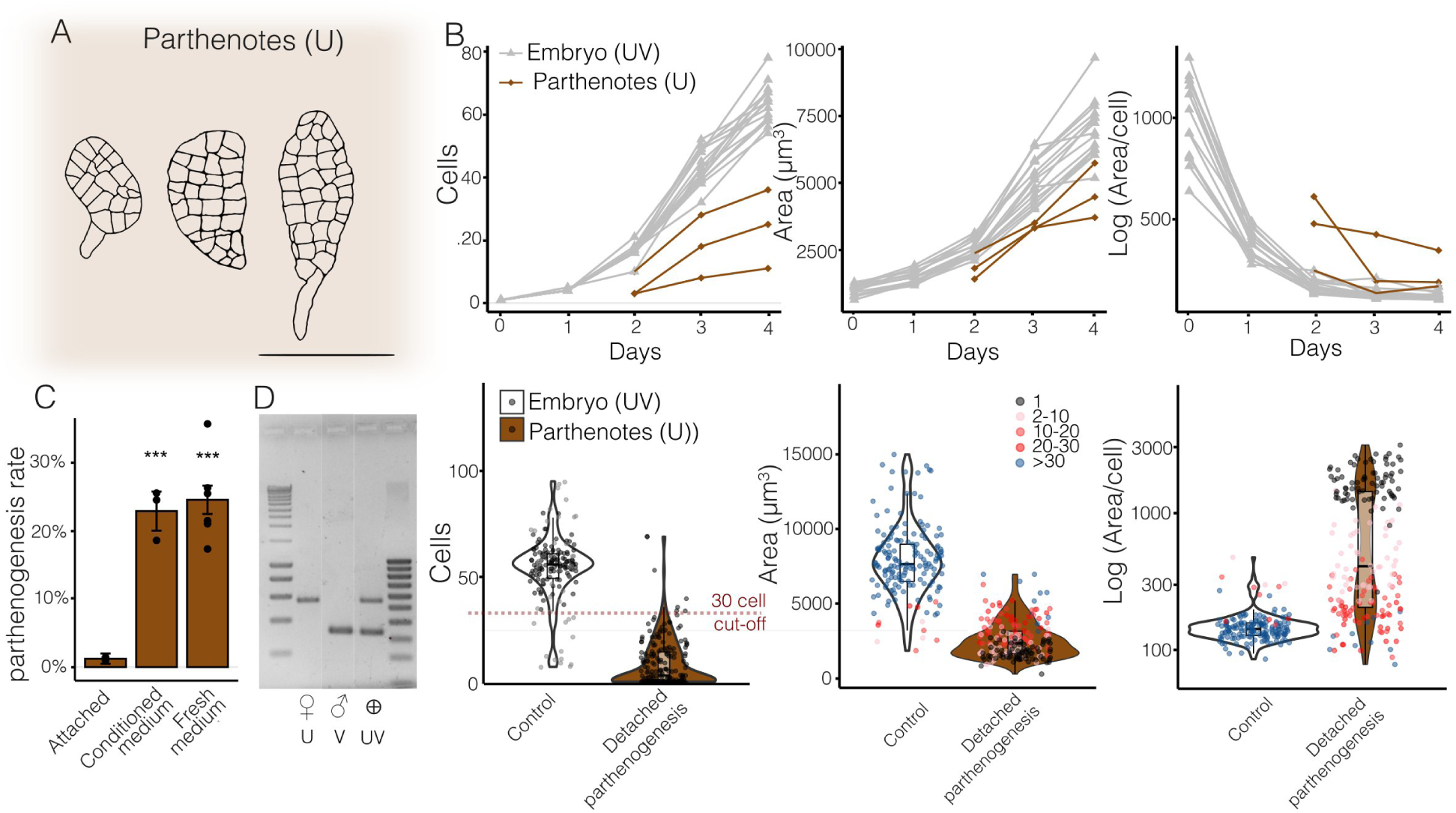
Parthenogenesis can be recognized by a slower growth and cell division. **(A)** Representative morphology of parthenotes at day 4. Scale bar = 50 µm. **(B)** Upper pannels, Patterns of cell division, area and average cell size of rep resentative parthenotes (red) vs embryos (grey) in the first 4 days of development. Lower panels, cell numbers, area and average cell sizes of embryos and detached parthenotes at 4 days after fertilisation (embryos) or detachment (parthenotes). The number of cells in parthenotes and embryos at day 4 is represented by colour: black, 1 celled; red, < 30 cells; blue > 30 cells. **(C)**. Parthenogenesis rate is significantly increased regardless whether fresh or female conditioned medium was applied (post hoc pairwise comparisons; ***, P value < 0.001). **(D)** Sex markers discriminate between female gametophytes or parthenotes (U), male gametophytes (V) and sporophytes (UV).

We compared growth kinetics of female parthenotes and embryos over four days (**Figure 3B**). Parthenotes exhibited a slower rate of cell division compared to embryos, rarely reaching the 30-cell state at day 4. In contrast, most embryos reached > 50 cells at day 4 (t = 31.058, df = 296.68, p-value < 2.2e-16; Welch Two Sample t-test, **Figure 3C**). Embryos were also significantly larger than parthenotes (t = 25.273, df = 229.33, p-value < 2.2e-16; Welch Two Sample t-test). The reduced size of parthenotes was due to a delay in the onset of the first cell divisions (**Figure 3B**). Indeed, while embryos showed a decrease in cell size during the first 2 days due to segmentation of the large zygote into smaller cells, in many of the parthenotes cell division did not keep up with growth, resulting in larger cells. Cell sizes in parthenotes were typically larger but showed an increased variance, reflecting a developmental delay. Only a small fraction (8%) of parthenotes reached the 30 cell-stage by day 4 (**Figure 3B**). Thus, the bulk of parthenotes can be clearly differentiated from embryos based on the cell number at day 4.

In *Ectocarpus*, parthenogenesis of unfertilized female gametes involves endoreduplication (Lewis et al. 1993, Bothwell et al. 2010). We found no evidence for endoreduplication in any of the analysed *Undaria* parthenotes (**Figure S4**). In order to further differentiate embryos from parthenotes, we used sex-specific PCR primers to genotype the detached embryos (**Figure 3C**). This allowed us to confirm the presence of male and female markers in detached embryos and exclude any potential enrichment in parthenogenetic embryos in our detached fraction.

Together, our results indicate that detachment from the maternal gametophyte induces parthenogenetic development that cannot be repressed by the presence of a gametophyte-conditioned medium. Parthenogenesis may be a confounding factor during analysis of early development in *Undaria*, but by assessing the number of cells at 4 days after germination and using PCR-based sex markers, we could effectively discriminate embryos from parthenotes.

### Attachment to the maternal tissue is required for robust apical basal patterning of the embryo

In order to test whether attachment of the egg to the maternal oogonium plays a role in early embryogenesis, we detached zygotes from the female gametophyte at the single-celled stage, and performed detailed morphometric measurements to follow the early development of the embryo. To avoid confounding with parthenotes, we identified embryos based on a combination of cell number (30-cell cutoff) and genotyping sex-markers.

A range of morphometric parameters were applied (*roundness*, *circularity* and *aspect ratio* (AR) to the 4-day old embryos (**Fig 4C**) and the effect of detachment was assessed using GLMMs. Overall, detached embryos were significantly rounder, more circular and had a lower AR compared to control embryos developing in contact with the maternal gametophyte (**Fig 4A, 4B, Table S1**). Because growth is expected to affect the shape of embryos, we included area in the GLM model. Inclusion of the area of embryos in the model did not result in a significantly better fit nor did it affect the significance of the effect of detachment (See Table S1 for GLMMs). Together, these analyses confirmed that detachment explained the difference in morphometric parameters better than individual size difference between embryos.

**Figure 4.**
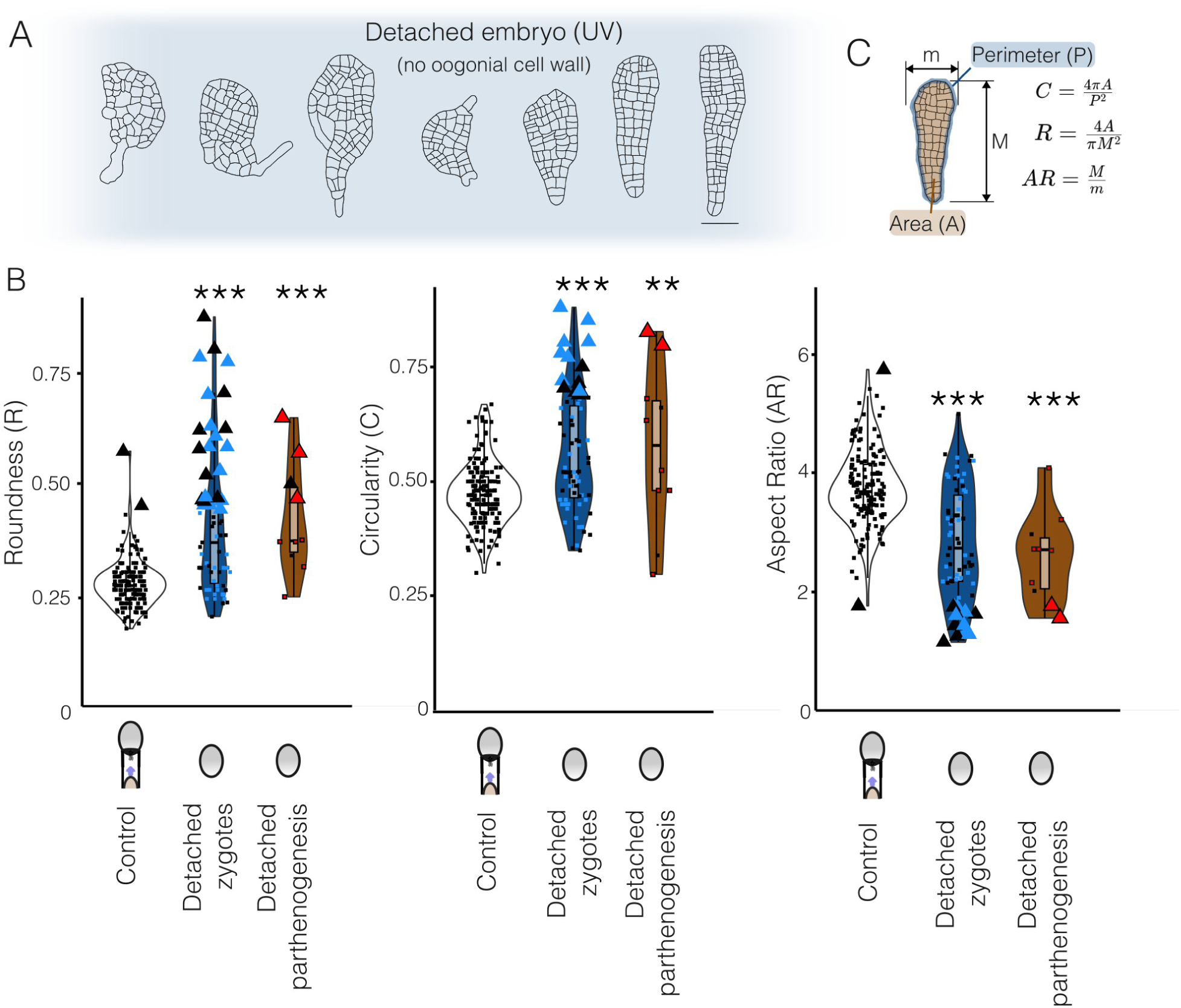
Detachment from the gametophytic oogonial cell wall affects robustness of the apical basal direction of the embryo leading to rounder embryos. **(A)**. Representative detached embryos with aberrant (left) or normal (right) apical basal patterning. **(B)** Morpho metric parameters for the synchronized control embryos; cut oogonial cell wall; detached zygotes (without oogonial cell wall) and detached parthenogenetic germlings. PCR confirmed crossed embryos are marked in blue, unconfirmed embryos in black, red sym bols denote presence of only female marker. Measurements that are outliers based on the control distribution (|z score| > 3) are high lighted with a larger triangles. Asterisk denote significant differences with the control (*** P value < 0.001; ** P value < 0.01, Supplemental table S1). **(C)** Different morphometric parameters used in this study C, circularity; R, Roundness; AR, Aspect Ratio.

The variation of detached phenotypes was larger compared to normal attached phenotypes. Respectively 34%, 20% and 18% of detached embryos qualify as outliers (|z-score| > 3) for roundness, circularity and aspect ratio measures compared to the control population (**Figure 4B**). Interestingly, these outlier fractions are also present among the few remaining parthenotes that developed more than 30 cells at day 4 (respectively 4, 2 and 2 outliers vs control distribution; N=10; Figure 4B). This suggests that detachment of both embryos and parthenotes results in a similar ratio of normal and aberrant phenotypes..

Amplification of male and female sex markers in all tested embryos ruled out confounding effects of parthenotes (36, N=76), potentially enriched in the detached fraction. This was further corroborated by the bimodal distribution in cell counts suggesting a clear-cut difference between the slowly developing fraction (suspected parthenotes) and the fast developing fraction (embryos) (**Figure S5**). Moreover growth parameters (number of cells or area) were not affected when putative parthenotes (< 30 cells at day 4) were omitted from the data (**Figure S4**). Because none of the single-celled embryos resulted in successful PCR amplification we cannot confirm their parthenogenetic nature.

### Developmental defects in detached embryos are rescued by the oogonial wall

To discern the cause of the embryo patterning defects, we tested whether the maternal cell wall provides a signal to the developing embryo. To this end, we detached the embryos from live maternal gametophyte cells by cutting the oogonial cell wall. A remnant part of the oogonial cell wall, however, remained attached to the embryo **(Figure 5A,B**). When the oogonial cell wall was present, the defect in apical-basal pattern was rescued. Cutting of the oogonial cell wall did not result in significantly different values for morphometric parameters in any of the GLMs (**Supplemental Table 1, Figure 5B**). Only detached embryos without a cell wall and detached parthenotes were morphologically different with significantly higher roundness, circularity and aspect ratio (**Figure 4B**, **Supplemental Table S1**). Our findings are most likely applicable to *Saccharina latissima* because a similar rescue effect of polarity defects could be seen by preserving the oogonial cell wall in a supplementary experiment (**Figure S6**), albeit without controlling for potential enrichment in parthenotes.

**Figure 5.**
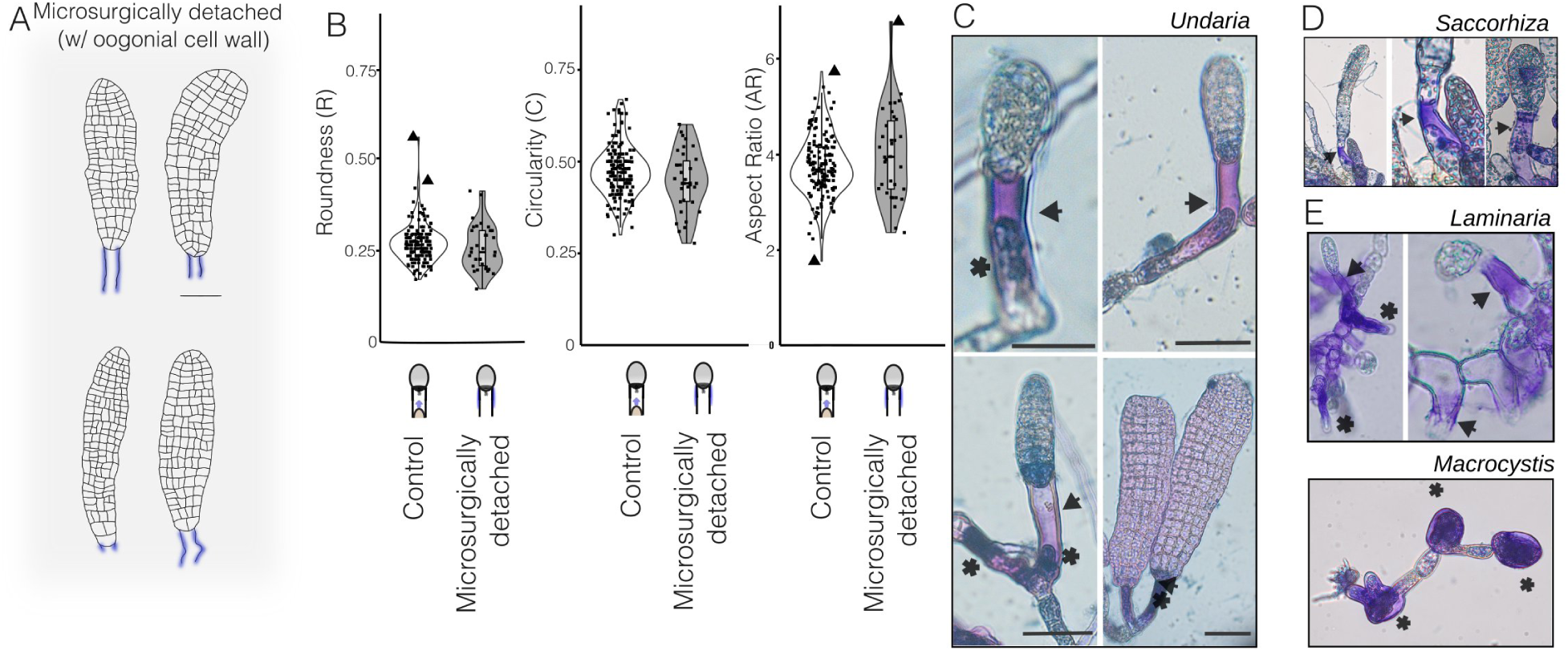
Preservation of the oogonial cell wall via microsurgical detachments rescues the phenotype and evolutionary conservation of deposition of Golgi derived F2 fucans assayed by TBO staining (at pH=1.5) as a marker for basal cell fate differentiation in the brown algae. **(A)** Representative embryos with oogonial cell wall marked in blue. Scale bar = 50 µm. **(B)** Rescue of apical basal patterning deffect if the oogonial cell wall is retained. Morphometric parameters did not differ significantly. **(C)** TBO in *Undaria pinnatifida* stains the oogonia, oogonial cell wall and sporophytic basal cells. **(D,E)**. Similar staining was observed in other kelp gametophytes such as those of (F) *Saccorhiza polyschides* (Tilopteridales) and (E) *Macrocystis pyrifera* (Laminariales) and *Laminaria digitata* (Laminariales) as in *Undaria pinnatifida*. Asterisks denote staining in oogonia and arrows denote TBO staining in oogonial cell wall bearing the sporophyte.

In the oogamous brown seaweeds *Fucus* and *Dictyota,* the apical-basal polarity is linked to localized cell wall modifications associated with the determination of the basal cell fate. Cell wall modifications can be visualized using Toluidine Blue O (TBO), which stains the sulfated F2 fucan fraction at pH 1.0-1.5 (Quatrano et al. 1976, Berger et al. 1994, Bogaert et al. 2017) (**Figure 5C**). In order to examine if the apical-basal polarity in *Undaria* is also associated with cell wall modifications, we used TBO on oogonial gametophyte cell wall and the attached embryos. Oogonial cell walls exhibited an intense staining before egg release (**Figure 5C**). After release, the oogonial cell wall still retained a more intense blue staining with TBO in comparison to the surrounding vegetative tissue (**Figure 5C**).

To test whether TBO staining (at pH 1.5) is conserved we stained other kelps including a member of the Tilpteridales (**Figure 5D**). A similar pattern was observed when staining cultures of Tilopteridales (*Saccorhiza polyschides*) as in *Undaria* (Laminariales) and other Laminariales (*Macrocystis pyrifera* and *Laminaria digitata*) showed similar staining pattern (**Figure 5D,E**). Therefore, the apical basal patterning of the sporophyte of kelps is associated with sulphatation of the cell wall of the oogonium (Quatrano et al. 1976), mirroring findings in other brown algal species *Dictyota* (Bogaert et al., 2017) and *Fucus* (Shaw & Quatrano et al., 1997) (**Figure 6**). While *Fucus* and *Dictyota* develop separated from the oogonium and show a deposition of sulfated fucans at the basal pole, kelps show a strong staining in the maternal oogonial cell wall (**Figure 5**).

**Figure 6.**
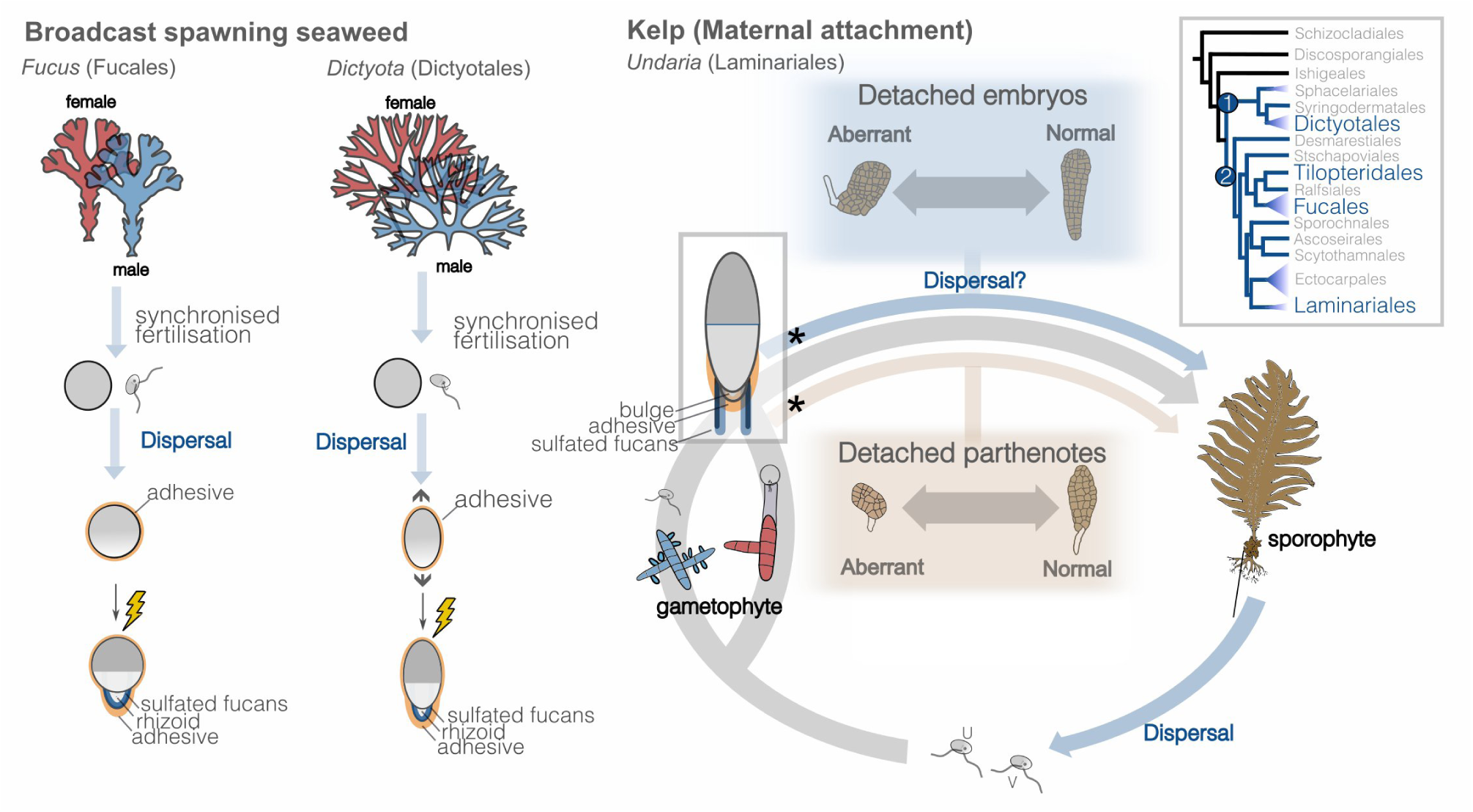
Development of broadcast spawners *Fucus* and *Dictyota* versus kelp with maternal attachment. **(left)**, *Fucus* and *Dictyota* release gametes synchronously in the surrounding medium producing large zygotes that will attach using an adhesive and will pho topolarize. During asymmetric cell division the basal cell wall is modified with sulfated fucans. **(right)** Effects of sporophyte attach ment and detachment in kelps. Simplified life cycle of kelps e.g. *Undaria* focussing on dispersal stages and parthenogenesis. Spores and dislodged sporophytes are seen as the dispersal stage in kelps, and sporophytes (or parthenotes) develop *in situ* on the gameto phyte. The here demonstrated viability of detached seedlings (parthenogenesis and embryo development) might contribute to dispersal of kelps (blue arrow). Blue and red boxes show variable developmental outcomes of detached embryonic and detached parthenogen etic developmental pathways. Life cycle can be be more complex in some species or strains of kelps (Ar Gall 1996, Shan & Pang 2021). Asterisks denote detachement of embryos or parthenotes. **(right, inset)** Evolutionary conservation of TBO basal staining in brown algae with SSDO clade (1) and BACR clade (2), containing the Laminariales and Tilopteridales.

## Discussion

Our results show that apical-basal polarity of embryo development in kelp is disturbed when the connection to the maternal gametophyte is disrupted and that preserving only the oogonial cell wall, enriched in sulfated fucans, is sufficient to rescue the defect in robustness of apical-basal patterning.

### Maternal guidance of apical basal patterning is mediated by the oogonial cell wall

Like many complex organisms, kelp embryos develop in close association with the maternal organism. In angiosperms the embryo is connected to the maternal organism via a suspensor and is influenced by signaling compounds and nutrients (Woudenberg et al., 2024). A role of the maternal tissue in buffering plant embryogenesis against environmental disturbances is suspected but not fully understood. Different degrees of contact to maternal tissue in non-seed plants are useful to disentangle this interaction (Woudenberg et al., 2024). The kelp maternal association represents a very loose association in which the embryo is not encapsulated as is the case for land plants. Our observations suggests that, nonetheless, the maternal gametophyte provides developmental robustness via the connection to the offspring sporophyte.

By providing one of the first comprehensive descriptions of early embryonic and parthenogenetic kelp development, we are able to compare developmental patterns across different brown algal model organisms. Our observations are in line with detection of the anchor flagella in *Saccharina* (Motomora 1991; Klochkova et al., 2019). The cell division pattern of *Undaria* embryos, with cells initially developing a unilayered 2-dimensional blade, are similar as described by Kimura et al. (2010), Klochkova et al. (2019) and Theodorou (2023). Like in *Arabidopsis* (Bayer et al., 2017) and *Dictyota* (Bogaert et al., 2017a, 2017b) the spherical egg cell undergoes an elongation along the apical-basal axis. Here, we show this elongation is an early process, occurring shortly after egg activation and therefore must have been completed for most of the zygotes at the time of detachment. Furthermore, the elongation is immediately accompanied by bulge formation. In *Dictyota* and *Fucus* (**Figure 6**), cell polarisation of the zygote is only completed late in the cell cycle allowing time to integrate environmental cues. Most likely, the elongation of *Undaria* zygotes depends on the maternal polarity in the polarised egg as very little time is left for environmental signaling. The elongation is, however, different from that of *Dictyota*, which elongates in a time-span of 90 seconds upon egg activation in a process involving shape change and no cell expansion (Bogaert et al. 2017b).

We show that detaching the zygote from the oogonial cell wall leads to patterning defects, but that a fraction of the detached zygotes are capable of forming seemingly well-patterned embryos. This pattern is mirrored in the developmental pattern of parthenotes of *Undaria* and in *Saccharina* (**Figu**r**e S6E, F**). We, therefore, conclude that the maternal connection provides developmental robustness to the embryo, but is not essential for its development.

The female gametophyte may make connection with the developing sporophyte. However, direct contact with the live maternal tissue – if restored at all – is only achieved late during embryogenesis and does not occur at all when the neighbouring cells differentiate also in an oogonium. The fact that retaining the oogonial cell wall is enough to rescue the defect in patterning robustness shows that direct interaction by the gametophyte via nutrients or hormones is unlikely to mediate the developmental robustness.

We hypothesize that the cell wall of the kelp oogonium is modified in a similar way as described for *Fucus* (Berger et al., 1994; Brownlee & Berger, 1995; Kropf et al., 1988; Quatrano & Shaw, 1997). In the latter, Golgi-derived vesicles (F-granules) deliver highly sulfated F2 fucans that can be stained using TBO (reviewed in Quatrano & Shaw, 1997). Interestingly, when rhizoid cells were ablated but the basal cell wall was maintained, this cell wall may direct cell fate causing differentiation into the basal cell fate (Berger et al., 1994). Here, we showed that, similar to *Dictyota* and *Fucus*, a similar positional correlation between sulfated fucans and basal cell identity occurs in two independently evolving clades of kelps (such as Laminariales and *Saccorhiza* (Tilopteridales)) (**Figure 5D,E**, **Figure 6**). This suggests that the mechanism that acts within the developing sporophyte of broadcast spawning algae has been co-opted by the female kelp gametophyte to aid basal cell differentiation in the sporophyte (**Figure 6**), therefore acting across life stages in kelps. We cannot exclude, however, that the cell wall may play an (additional) role by keeping the flagellar root in place or alternatively providing a physical constricting force enabling bulge formation.

### Partheno sporophytes can be discerned from embryos

A round morphology and a tendency to easily detach are two properties that are traditionally used to discern parthenotes from embryos during crossing experiments (tom Dieck et al., 1992; Martins et al., 2017; Liboureau et al., 2024). Parthenogenesis is marked by a higher likelihood of detachment (Klochkova et al., 2019; Martins et al., 2017), reduced growth, and various defects in mitosis and polarity leading to rounder embryos (Schreiber 1930; tom Dieck 1992, Ar Gall et al. 1996; Fang et al., 1979). While most parthenotes are not viable, a proportion of them may grow into reproductive adult seaweeds (Fang 1979; Lewis et al., 1993; Müller et al., 2019; Camus et al., 2021; Shan et al., 2021).These observations often co-occur with polyploidy, suggesting that similarly to *Ectocarpus* (Bothwell et al. 2010) endoreduplication may occur during parthenogenesis (Lewis et al., 1993; Shan et al. 2001). While the (largest) individuals we obtained in culture flasks were all haploid, we cannot exclude that diploidisation occurs in other parthenotes as reported for *Undaria* (Lewis et al. 1993, Shan et al. 2001).

Being correlated to both the treatment (detachment) and the outcome (abnormal development), parthenogenesis has to be controlled for as a potential confounding variable. The issue of parthenotes, is stressed by the observation that upon detachment a 20-fold increase of parthenogenetic development occurs in unfertilized eggs from female-only cultures (**Figure 3C**). Here we confirmed that in *Undaria* the detached fractions are not enriched in parthenogenetic germlings. While initial experiments were conducted in *Saccharina latissima* (**Figure S6**) a strong enrichment in parthenotes was suggested by the slower growth of the detached fraction. Therefore we resorted to an *Undaria* strain which shows much lower parthenogenesis rates.

### The kelp (partheno )sporophyte as a dispersal stage

Understanding the mechanisms of kelp dispersal is important to (i) estimate the degree of gene flow from commercially grown kelps to native populations, (ii) understand the ways of dispersal in kelp restoration projects (Vanderklift et al. 2020) and (iii) understand the dispersal vectors for non-native kelp such as *Undaria pinnatifida* (Hay et al. 1990). Kelps such as *Undaria* are thought to disperse via the zoospore stage (short-range) and via dislodged fertile sporophytes (long-range) (Forrest et al. 2000, Vanderklift et al. 2020, Edwards 2022). Due to the physical connection to the maternal gametophyte, the zygote and embryo are usually not considered as dispersal vectors. The observation that embryos may lead to individuals bearing rhizoids (Figure 4A), despite significant patterning defects, raises the question to what extent detached development is relevant in the field and contributes to kelp dispersal in a similar fashion as is the case for broadcast spawners (**Figure 6B**).

Additionally, detachment of unfertilised eggs from the gametophyte triggers parthenogenetic development. Because at least some parthenotes of kelp species, including *U. pinnatifida*, have been shown to acquire a normal morphology and even to become fertile (Fang 1979; Lewis et al., 1993; Müller et al., 2019; Camus et al., 2021; Shan et al., 2021), detachment of parthenotes and embryos by for example wave action, grazing or swaying seaweeds may provide an additional dispersal stage mediating gene flow, in addition to released meiospores and rafting of dislodged fertile sporophytes. For the case of parthenotes, field studies show that parthenogenetic kelp sporophytes are present in natural populations and may even reproduce, producing only female gametophyte offspring (Oppliger et al. 2007; Klochkova et al., 2017). However while fully parthenogenetic range edge populations are described for other brown algae (Hoshino et al., 2019; 2021), population genetic studies of *Undaria* have as yet not reported any signals of widespread clonality (Shan et al., 2017; Guzinskiet et al., 2018, Shan & Pang 2022), indicating that the role of parthenogenesis in these populations is not a dominant factor.

### Bringing the kelp model up to par with broadcast spawning seaweeds

Besides relevance for developmental biology of kelps, our findings are also relevant from a methodological perspective. It is important to note that detached embryo development is often still normal and without significant delay unlike parthenogenesis. Therefore detached development might still be considered for some applications such as automated phenotyping, as long as the polarity defects are taken into account or irrelevant to the assay (e.g. individual photosynthetic rates; compound production).

In broadcast spawners like *Fucus* or *Dictyota*, it is easy to obtain large populations of synchronously developing zygotes free from viable parthenotes and maternal tissue, making them suitable models for studying the installation of apical-basal polarity and the asymmetric cell division of the zygote (Bogaert et al., 2022; Coelho & Cock, 2020). In kelps however, the physical attachment poses challenges, resulting in mixtures of different day cohorts (Theodorou et al., 2021). Here we mitigated the negative implications of primary attachment on fertilization and polarization assays. First, we detailed a protocol to navigate these challenges, enabling the production of synchronized zygote batches (see methods). While kelp fertilisation is synchronous by nature with egg release at the onset of darkness (Lüning, 1981), the attachment of eggs to the maternal gametophyte and the repeated gamete release over days obscures potential recruitment cohorts. Our approach allowed us to produce one synchronized batch that remained attached to the oogonia by adding only male gametes. Secondly, we examined the nature of detached zygote development compared to normal (attached) development, to evaluate whether and to what extent maternal signaling contributes to sporophyte development. Thirdly, we had to address the issue of parthenogenesis. Here we showed that parthenogenetic germlings can be recognized by a growth delay and that populations of detached zygotes can be validated by genotyping using sex markers. By mitigating these three drawbacks, we bring kelp embryos up to par with more established models for the study of early development.

### Conclusions

In conclusion, we showed that apical-basal polarity in the kelp embryo is less robust when detached, resulting in defects in apical-basal patterning for a significant part of the population. We show that the robustness of apical-basal polarity is restored when the cell wall remains attached, suggesting a vital role for the cell wall of the oocyte connection. We further assessed the hypothesis that the robust apical-basal patterning is influenced by the deposition of Golgi-derived cell wall material by TBO staining, a method previously used to demonstrate the establishment of basal cell fate in brown algae as in the case for *Fucus* (Shaw & Quatrano 1996, Quatrano & Shaw 1997). The here proposed working hypothesis suggests that the fate of the basal cell is determined by substances secreted into the oogonial cell wall by the maternal gametophyte reminiscent to the mechanism in the brown alga *Fucus* (Berger et al., 1994; Kropf et al., 1988). These positional cues contribute to the robustness of developmental patterning, ensuring that the emerging sporophyte develops a defined orientation and structure.

## Acknowledgements

We thank Sofie D’Hondt, Francesca Petrucci, Laura Strickx for assistance with dissections, genotyping and culture facilities. We thank Tijs Ketelaar and Jessica Knoop for stimulating conversations. We thank Luna van der Loos and Bertrand Jacquemin for sampling *Undaria* and *Saccharina,* respectively. This work was supported by the Max Planck Society, the ERC (grant n. 864038 to SMC), Flanders Innovation & Entrepreneurship-Interdisciplinair Cooperatief Onderzoek (VLAIO-ICON) project (Blue Marine.com3, AIOSBO2019001503), EMBRC Belgium and the Research Foundation Flanders (projects I001621N).

## Author contributions

ED: Investigation (lead), Methodology (equal), Visualization (lead), Writing – original draft (supporting), Writing – review and editing (supporting). YM, FG, JB, EZ: Investigation (supporting), Methodology (equal), Writing – review and editing (supporting). DL: Investigation (supporting), Writing – review and editing (lead). TB: Conceptualization (supporting), Writing – review and editing (supporting). SMC, ODC: Conceptualization (lead); Funding acquisition (lead); Supervision (equal); Methodology (equal); Writing – review and editing (lead). KB: Conceptualization (lead); Investigation (supporting); Supervision (equal); Methodology (equal); Visualization (lead), Writing – original draft (lead); Writing – review and editing (lead).

## Supplemental figures

**Figure S1.**
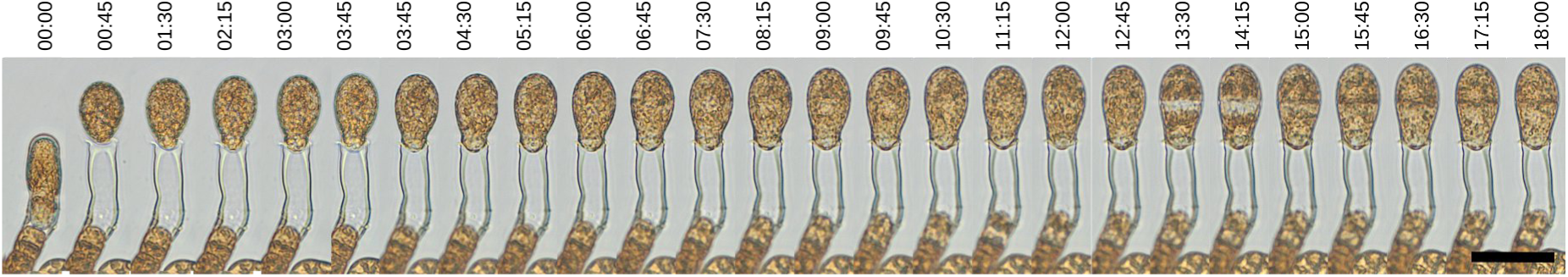
Time lapse 1. Developing zygote undergoing fast elongation (expansion) along the apical basal polarity vector and asymmetric cell division.

**Figure S2.**
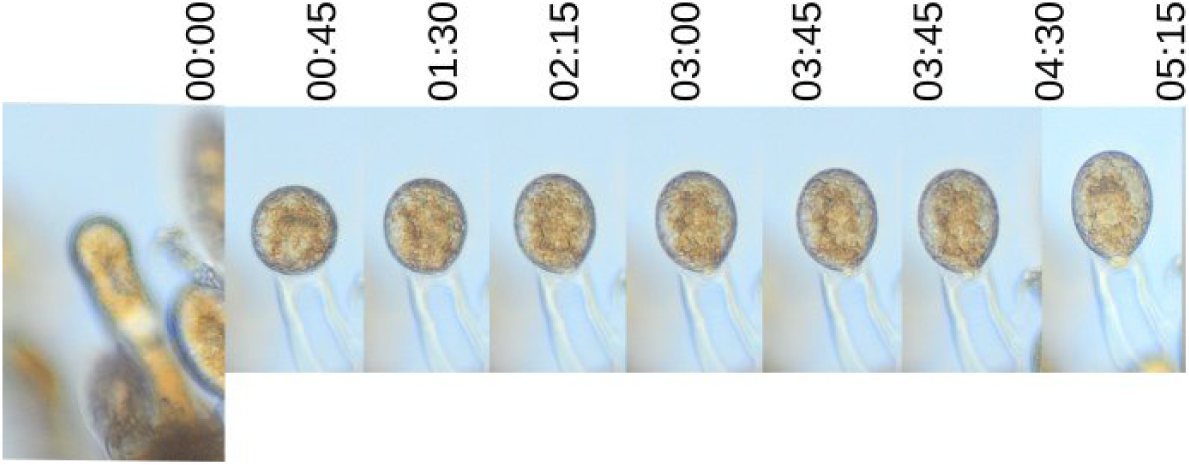
Time lapse 2. Developing zygote undergoing elongation (expansion) along the apical basal polarity vector.

**Figure S3.**
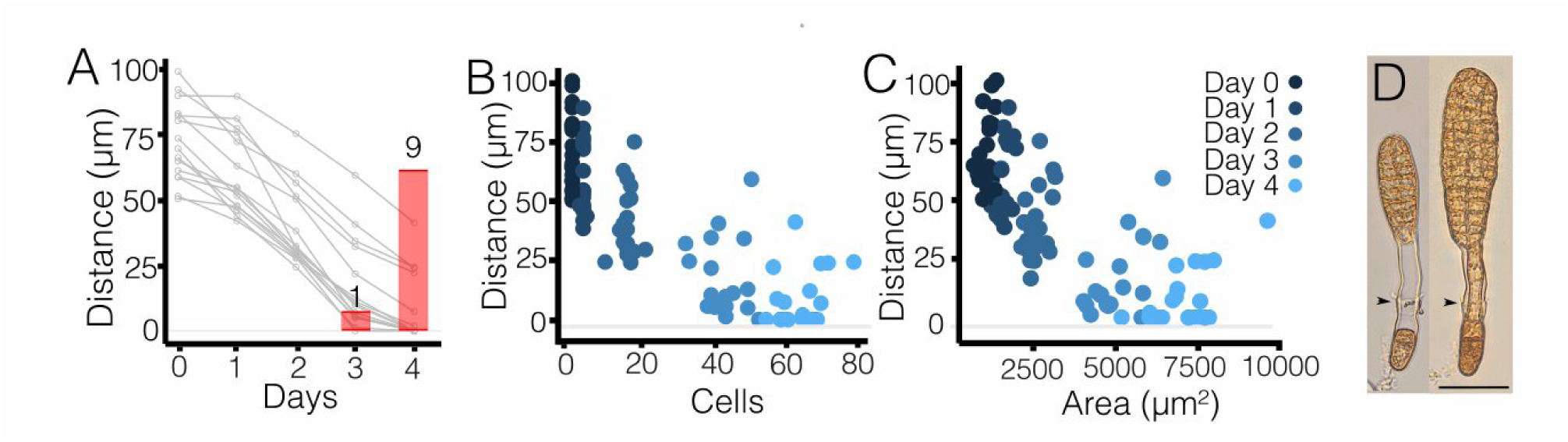
Sporophyte develops during first 3 4 days only in direct contact with the oogonial cell wall of the female. **(A)** Female gametophyte filament makes contact with the sporophyte only around 4 days AF. Red bars denote the number of time lapses followed over 4 days AF (N=14). (**B, C)** Distance of the developing embryo to the nearest living gametophyte cell as a function of the number of cells (B) and area (C) of the developing embryo. Blue shades denote real time AF.

**Figure S4.**
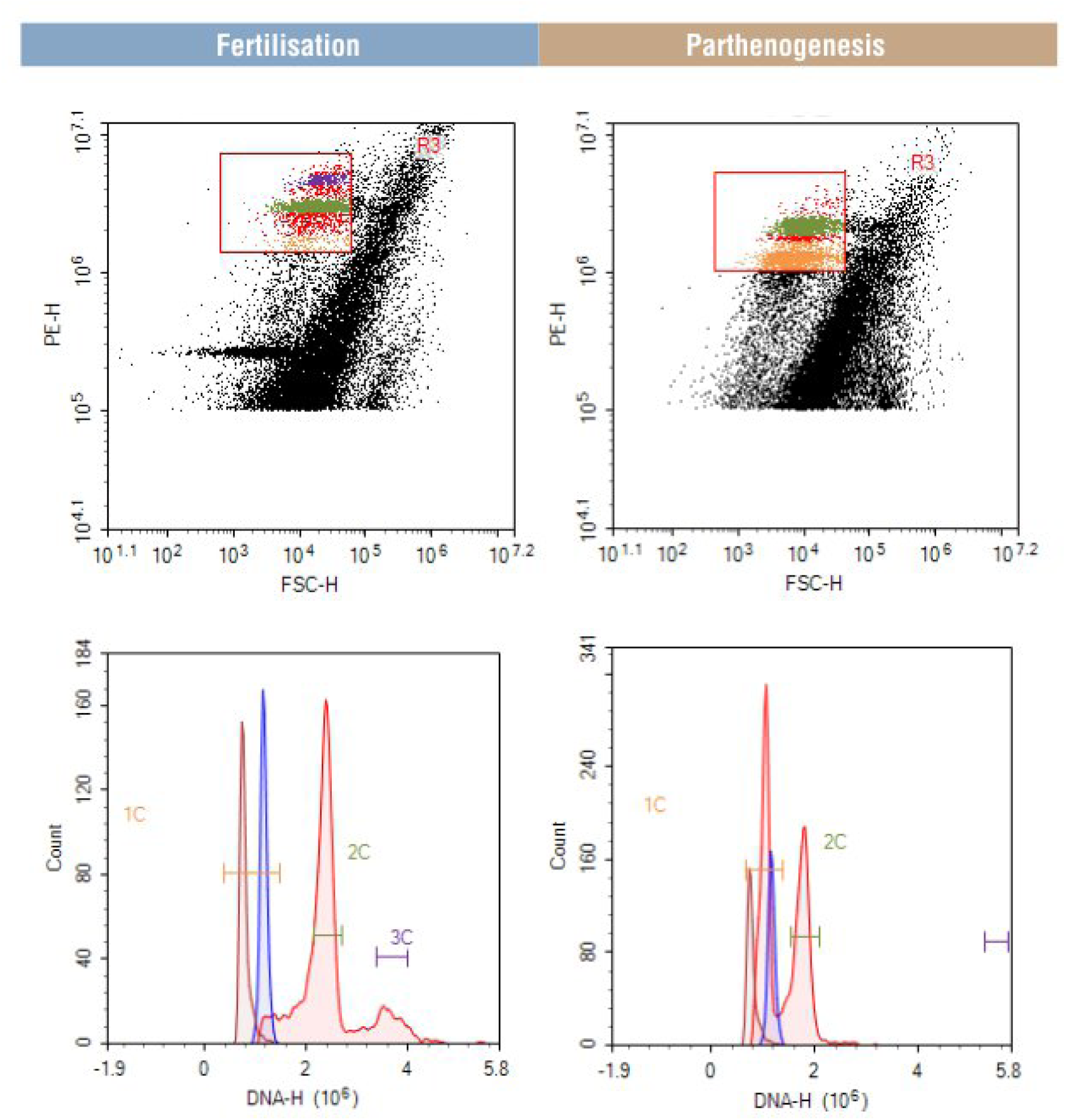
Results of flow cytometry analysis to estimate DNA content of *Undaria pinnatifida* sporophyte strains from one fertilized (control) population and one parthenogenetic individual. Plots in the top row show populations of nuclei grouped by DNA content in scatter plots. Bottom row shows density plots of fluorescence signals as a proxy for DNA content. Brown and blue peak inserts refer to measurements of haploid gametophytes; red peaks represent the tested (partheno)sporophyte nuclei.

**Figure S5.**
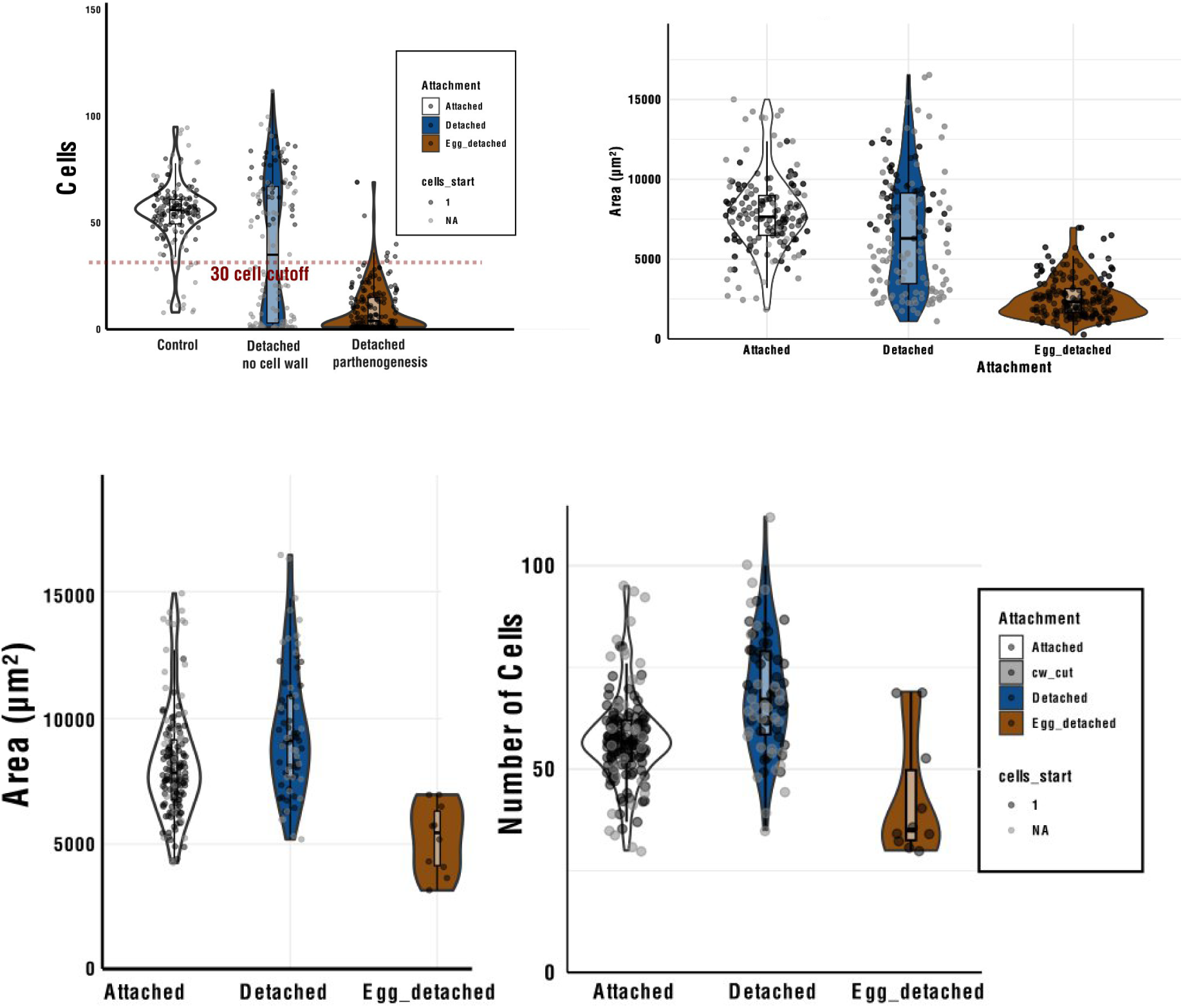
Slow cell division as a marker for parthenogenesis vs normal development (detached or attached.) No decrease in cell division could be observed.

**Figure S6.**
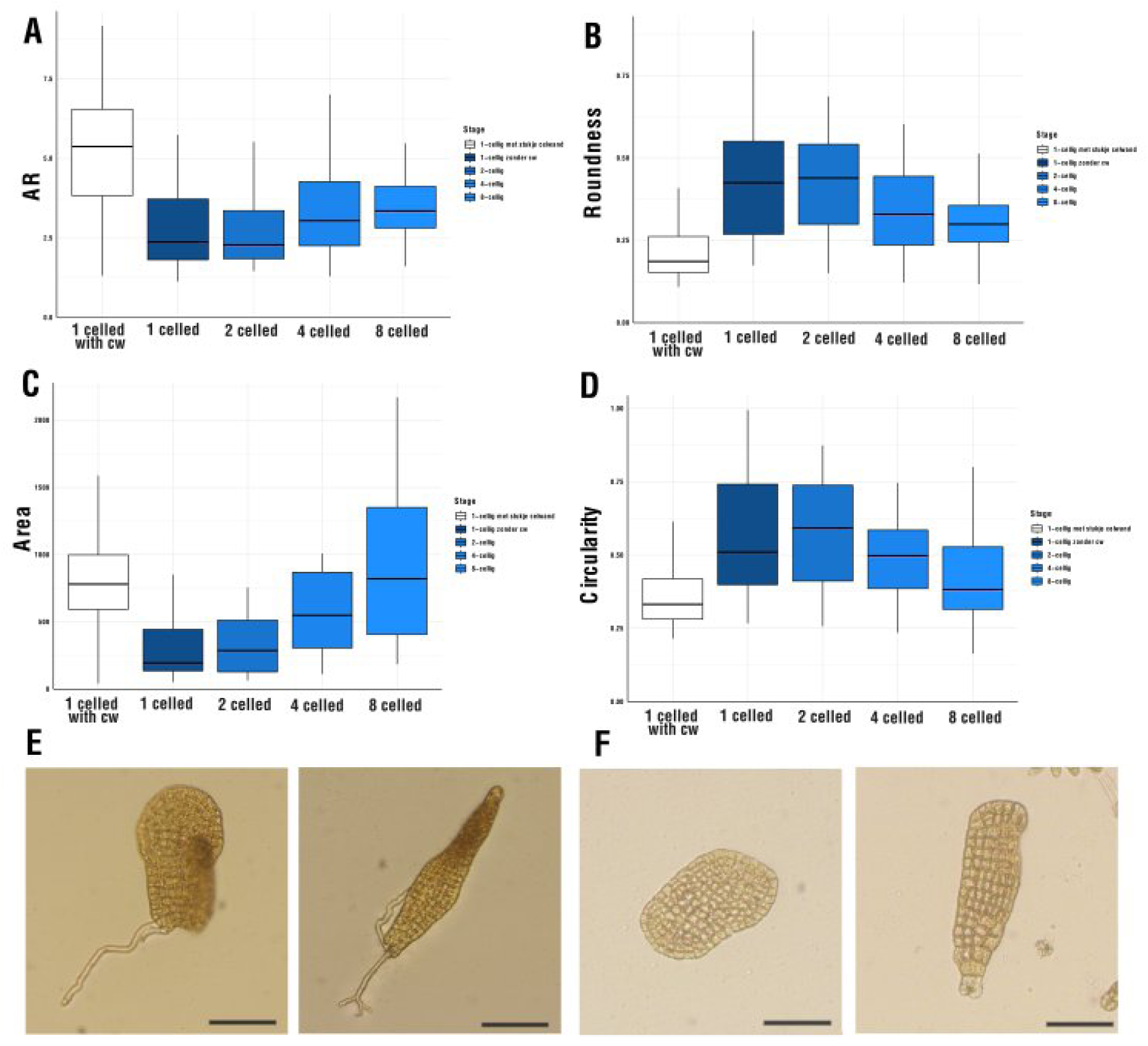
Rescue effect of the cell wall can also be observed in *S. latissima*. (**A,B,D**) In contrast to *U. pinnatifida*, 2,4 and 8 celled embryos could be easily detached. When detached in a later stage the effect of detachement gets lower as expected because the embryos could develop with maternal guidance during the early stages. 1 celled cells show the strongest circularity effect. In this preliminary experiment however, growth was affected as well **(C)**, suggesting parthenogenesis could be a confounder and therefore this preliminary experiment should be interpreted with care. **(E)**, when detached putative zygotes develop both aberrant (rounder) blades (left), or more normal and elongated (right). **(F)**, parthenogenetic detached development resembles normal detached development. Scale bars = 50µm.

**Table S1.**
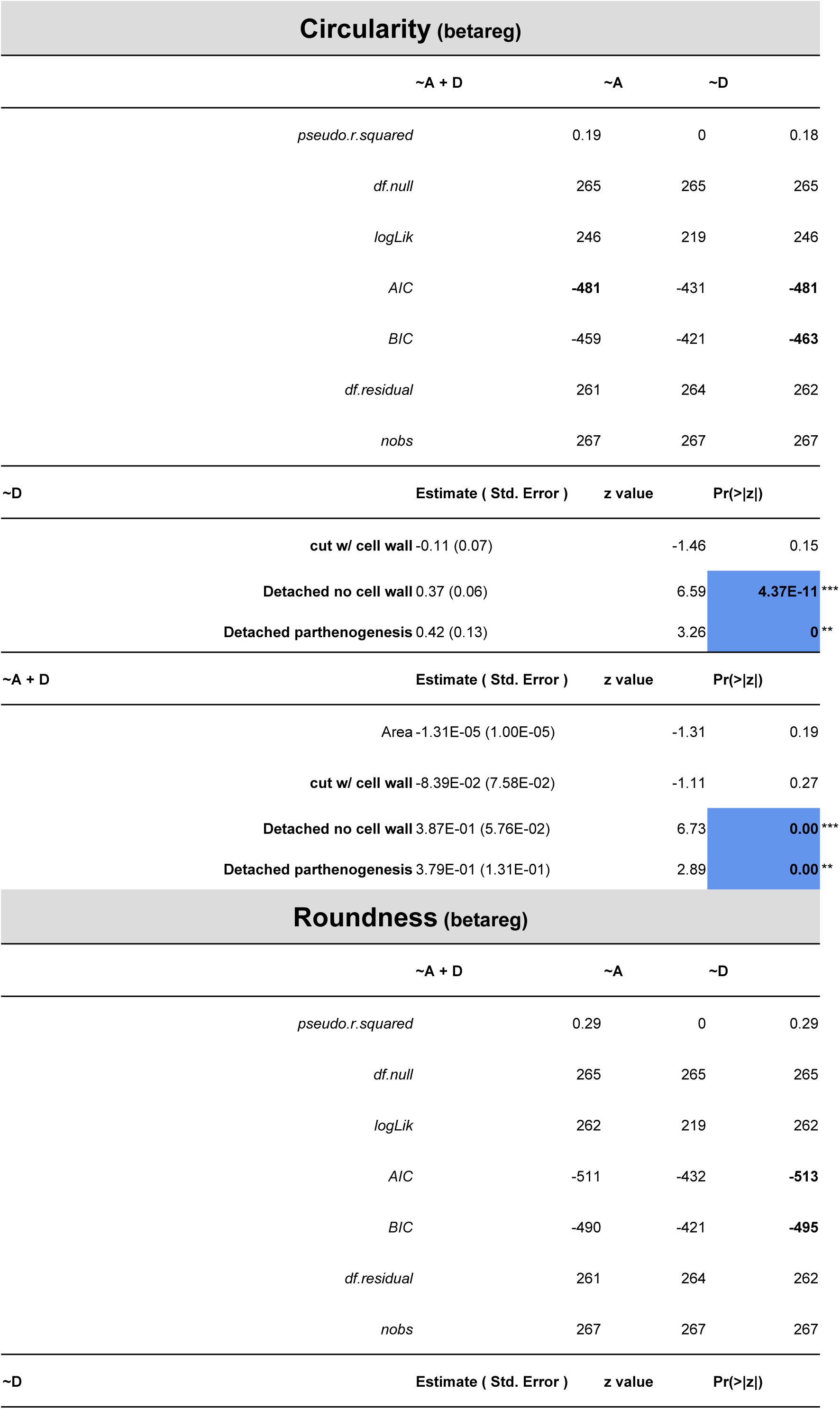
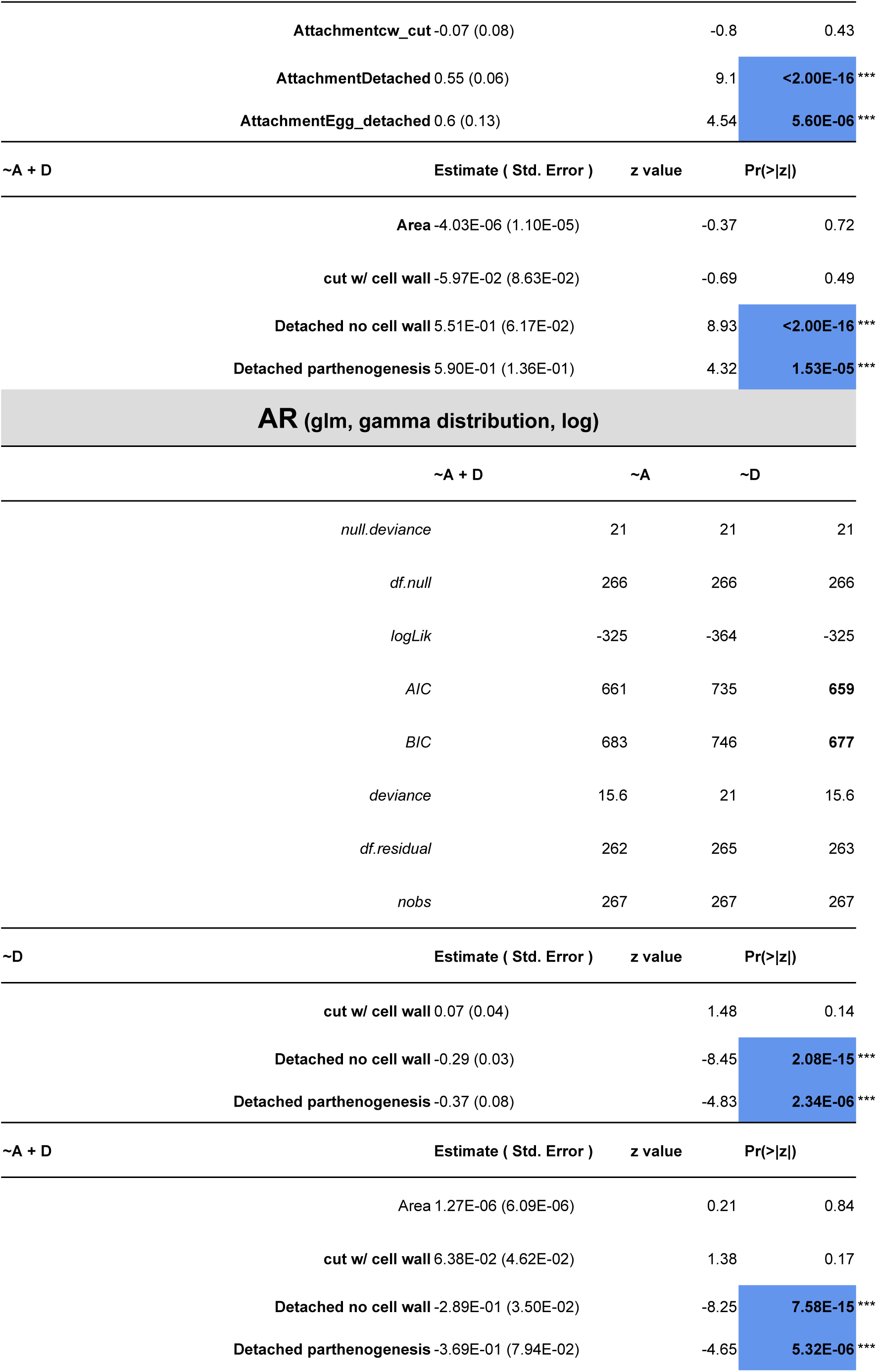
Table describing results from GLM modelling respectively circularity, roundness and AR in response to growth (Area, A) and detachement (synchronised control, cut w/cell wall, detached parthenogenesis) at 4 days after fertilisation (or detachement).

